# Dynamics of functional networks for syllable and word-level processing

**DOI:** 10.1101/584375

**Authors:** J.M. Rimmele, Y. Sun, G. Michalareas, O. Ghitza, D. Poeppel

## Abstract

Speech comprehension requires the ability to temporally segment the acoustic input for higher-level linguistic analysis. Oscillation-based approaches suggest that low-frequency auditory cortex oscillations track syllable-sized acoustic information and therefore emphasize the relevance of syllabic-level processing for speech segmentation. Most linguistic approaches, however, focus on mapping from acoustic-phonemic representations to the lexical level. How syllabic processing interacts with higher levels of speech processing, beyond segmentation, including the anatomical and neurophysiological characteristics of the networks involved, is debated. Here we investigate the effects of lexical processing and the interactions with (acoustic) syllable processing by examining MEG data recorded in two experiments using a frequency-tagging paradigm. Participants listened to disyllabic words presented at a rate of 4 syllables/sec. Two conjectures were evaluated: (i) lexical processing of words activates a network that interacts with syllable processing; and (ii) syllable transitions contribute to word-level processing. We show that lexical content activated a left-lateralized frontal and superior and middle temporal network and increased the interaction between left middle temporal areas and auditory cortex (phase-phase coupling). Mere syllable-transition information, in contrast, activated a bilateral superior-, middle temporal and inferior frontal network and increased the interaction between those areas. Word and syllable processing interacted in superior and middle temporal areas (cross-frequency coupling), whereas syllable tracking (cerebro-acoustic coherence) decreased when word-level information was present. The data provide a new perspective on speech comprehension by demonstrating a contribution of an acoustic-syllabic to lexical processing route.

**Significance statement:** The comprehension of speech requires integrating information at multiple time scales, including phonemic, syllabic, and word scales. Typically, we think of decoding speech in the service of recognizing words as a process that maps from phonemic units to words. Recent neurophysiological evidence, however, has highlighted the relevance of syllable-sized chunks for segmenting speech. Is there more to recognizing spoken language? We provide neural evidence for brain network dynamics that support an interaction of lexical with syllable-level processing. We identify cortical networks that differ depending on whether lexical-semantic information versus low-level syllable-transition information is processed. Word- and syllable-level processing interact within MTG and STG. The data enrich our understanding of comprehension by implicating a mapping from syllabic to lexical representations.

## Introduction

Oscillation-based approaches to speech comprehension posit that temporally segmenting the continuous input signal (Ahissar and Ahissar, 2005; Ghitza and Greenberg, 2009; Panzeri et al., 2010; Ghitza, 2011; Giraud and Poeppel, 2012; Zion Golumbic et al., 2012) is realized through phase-alignment of low-frequency (< 8 Hz; delta-theta) neuronal oscillations in cortex to the slow fluctuations of the speech signal at the syllabic scale (Poeppel, 2003; Luo and Poeppel, 2007; Gross et al., 2013; Fontolan et al., 2014; Doelling et al., 2014; Rimmele et al., 2015). This imposes constraints (Rimmele et al., 2018b) such that speech perception is optimal at syllabic rates that fall within the range of intrinsic auditory cortex oscillations in the delta-theta range (Ghitza and Greenberg, 2009; Ghitza, 2014, 2016, Teng et al., 2017).

Several experiments have documented hierarchical processing over linguistic processing levels, such as words, phrases, and sentences (Ding et al., 2016; Martin and Doumas, 2017; Molinaro and Lizarazu, 2018; for review Meyer, 2017; Rimmele et al., 2018a). Different types of evidence point to interactions of higher level processing with syllable-level segmentation. For example, predictions arising from multiple levels, e.g. phonological or syntactic/semantic processing (Marslen-Wilson and Tyler, 1980; Altmann and Kamide, 1999; Kotz and Schmidt-Kassow, 2015; Scontras et al., 2015; Jadoul et al., 2016), modulate lower levels. The top-down modulation of phase-alignment of oscillations to speech acoustics is suggested by increased speech-tracking for intelligible speech (Peelle et al., 2013; Zion Golumbic et al., 2013; Rimmele et al., 2015; Park et al., 2015), which is accompanied by increased connectivity between auditory cortex and higher-level processing areas, including frontal and motor cortex (Park et al., 2015, 2018). At what linguistic levels the processes interact (Keitel et al., 2018) and whether this is due to lexical-semantic (Peelle, 2012; Peelle and Davis, 2012) or phonological processing (Di Liberto et al., 2015; Mai et al., 2016) is debated. Puzzles remain, both at the neural and the linguistic/psycholinguistic levels. Regarding the former, the spectral characteristics of the network dynamics are largely unknown. Delta-theta neuronal oscillations are crucial for tracking syllable-sized acoustic chunks; however, whether (i) the areas implicated in syllable processing interact with higher-level processes through phase-synchronization at these frequencies (Park et al., 2015; Assaneo and Poeppel, 2018) or through cross-frequency coupling (Keitel et al., 2018), (ii) networks involved in higher-level processing synchronize in a similar spectral range, and (iii) whether the type of connections are feedback or feedforward (Fontolan et al., 2014; Fries, 2015; Michalareas et al., 2016), merit study.

The posterior MTG (left lateralized, Binder et al., 2009) provides a sound-to-meaning interface, mapping phonological to lexical representations (Hickok and Poeppel, 2000, 2004; Binder et al., 2009; Gow, 2012; Rodd et al., 2015). *Sublexical* contingencies, such as syllable transitions, contribute to word processing, and have been shown to activate parts of the STS (Okada and Hickok, 2006; Mesgarani et al., 2014) and a dorsal-path network (Hickok and Poeppel, 2007). Additionally, parts of the IFG have been shown to contribute to word-level processing, e.g. in sublexical processing tasks reflecting sensory-motor integration (Hickok and Poeppel, 2007) or in tasks that elicit lexical competition (Thompson-Schill et al., 1997; Kan et al., 2006; Rodd et al., 2015).

Using a frequency-tagging paradigm (Buiatti et al., 2009; Ding et al., 2016) to ‘align’ the neuronal processing of syllables and words, we address the following: do lexical and syllable-transition cues of words activate a network which is left lateralized for lexical processing? And, crucially, do (acoustic) syllable- and word-level processing interact? In two MEG experiments, native German speakers listened to isochronously presented syllables of a native and a foreign language, presented at 4 syllables/sec, resulting in a rate of 2 Hz for ‘disyllabic units’, i.e. words or pseudowords. The effects of lexical information (native vs. foreign condition), lexical plus syllable transition information (native vs. foreign pseudo-words), and of syllable transition information alone (foreign vs. foreign pseudo-words) and the interaction of these processes with syllable processing are characterized.

## Materials and Methods

### Participants

German native speakers with no previous knowledge of Turkish participated in the two MEG experiments. In the first experiment the data of 18 (mean age: 24.32 years; SD: 3.30 years; female: 10) healthy right handed (Oldfield mean score: 91.38, SD: 15.44) participants are included in the analysis (right hand “yes” button: n=9). In the second experiment the data of 19 (mean age: 24.46 years; SD: 3.73 years; female: 10) healthy right handed (Oldfield mean score: 92.74, SD: 14.66) participants are included in the analysis (right hand “yes” button: n = 10). Several participants were excluded: because of outlier behavioral performance (accuracy < mean – 2 *SD; Exp. 1: n = 2; Exp. 2: n = 1) and because of technical issues (no triggers, audio problems; Exp. 1: n=2; Exp. 2: n = 1). Individual T_1_-weighted MRI scans were conducted for all participants, except for some participants who either did not match the MRI criteria or did not show up to the MRI scan session (Exp. 1: n = 5; Exp. 2: n = 3). The study was approved by the local ethics committee of the University Hospital Frankfurt. All participants gave written informed consent for participating in the study and received monetary compensation.

### Experimental Design

#### Paradigm

Participants were asked to listen to sequences of disyllabic words, that contained either German, Turkish, or pseudo-words. In Exp. 1, sequences of German words (Fig. 1 B) and Turkish words (Fig. 1 C) were used. In Exp. 2, sequence of German words (Fig. 1 B) and Turkish pseudo-words (Fig. 1 D) were used. Each sequence contained 38 syllables that formed 19 ‘disyllabic units’ (word or pseudo-words). By presenting isochronous syllables, the presentation rate for syllables was fixed at 4 Hz, resulting in an occurrence of disyllabic units at 2 Hz. In order to maintain participants’ attention on the auditory stimuli, a target stimulus, which consisted of a disyllabic unit that was made of a repetition of the same syllable, was inserted in 28.57% of the trials. Each block contained 29% trials with a target stimulus (equally distributed across conditions). Participants were asked to indicate with a button press whether a target stimulus was present after each trial (Fig. 1 A; index finger left and right hand; the response hand was counterbalanced across participants). Each trial was followed by a jittered intertrial interval (2 - 3.5 sec).

**Figure 1.**
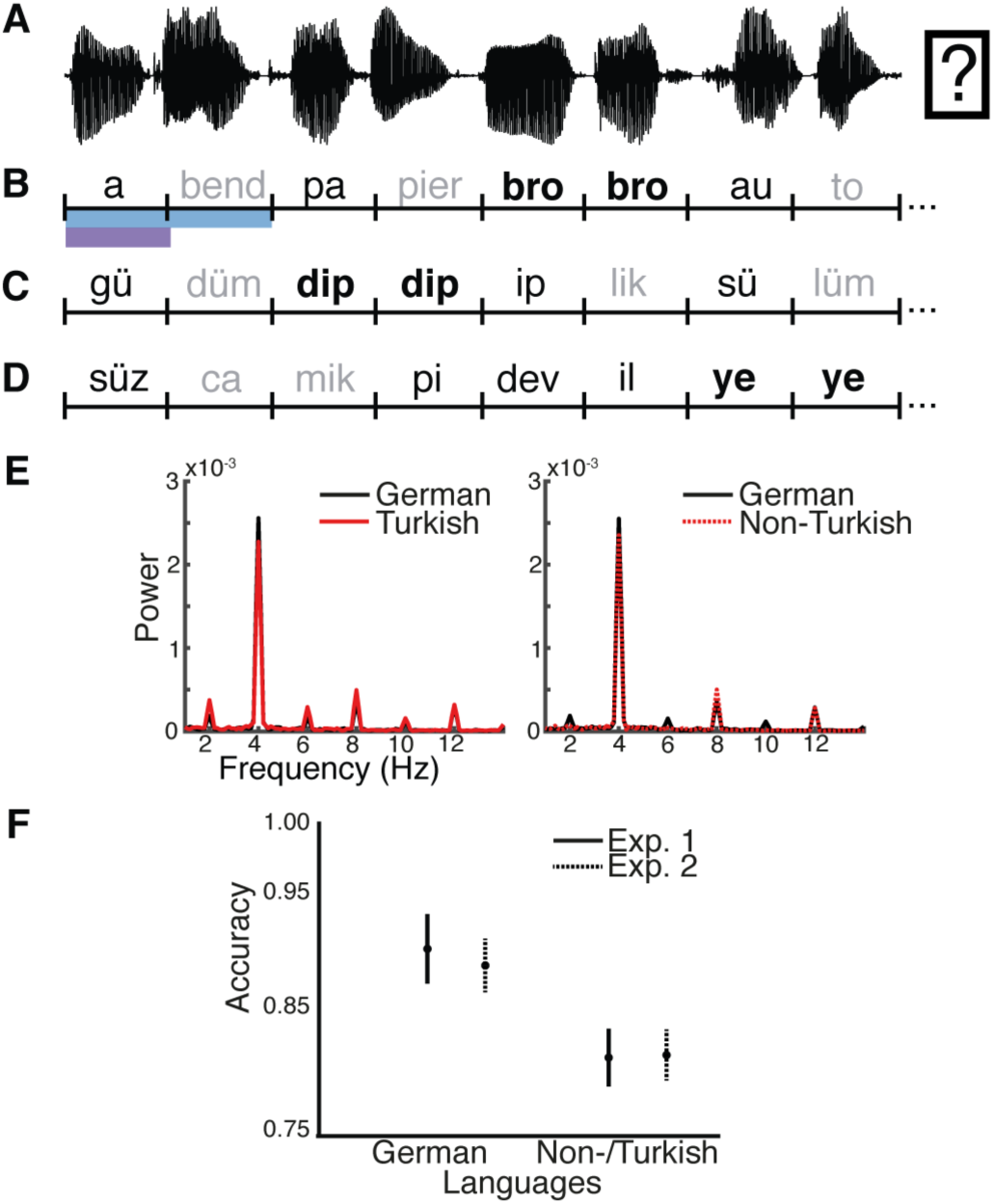
Schematic of the Paradigm and Material. (A) Illustrates the structure of a trial: participants indicated after listening to syllable sequences whether a target (bold) was present; In both experiments syllables were presented at a rate of 4 Hz (250 ms; purple bar); (B) In Exp. 1 in the German condition syllables can be grouped into disyllabic words (black: first syllable, gray: second syllable) at a rate of 2 Hz (500 ms; blue bar); (C) In Exp. 1, in the Turkish condition the syllables presented at 4 Hz cannot not be grouped into words based on semantic knowledge; In Exp. 2, the German condition corresponded to that of Exp. 1; (D) In Exp. 2, in the Non-Turkish condition syllables, presented at 4 Hz, cannot be grouped into words based on semantic knowledge or syllable transition information; (E) The speech acoustics show a strong peak at the frequency-tagging (syllable) rate (4 Hz) in Exp. 1 (left column) and Exp. 2 (right column); (F) Overall, the performance (accuracy) did not differ across experiments, however, overall was higher for the German compared to the Turkish/Non-Turkish conditions (error bars: +-1 SEM).

#### Stimulus Selection

German disyllabic words were selected from the CELEX lexical database (Baayen et al., 1995). In order to maximize the speed of lexical access of the word as well as the predictability of the second syllable within a word, we selected words with a high frequency (CELEX spoken word frequency: MannSMln ≥ 10; Celex zeros replaced by Leiziger Wortschatz Corpus; LWC ≥ 4000) and with a high transition probability between the two syllables (STP ≥ 0.3 %). The STP was calculated for all disyllabic lemmas in the corpus, by dividing the wordform frequency of each lemma by the sum of the wordform frequencies of each wordform that contained the first syllable of the token (Lewis et al., 2011). Laplace transformation of zero frequencies was used (Brysbaert and Diependaele, 2012). Turkish disyllabic words were selected from the TELL database (http://linguistics.berkeley.edu/TELL/; e.g. Scharinger et al., 2011) and manually checked by a Turkish native speaker (for wordness). In total, 134 German words and 134 Turkish words were selected (noun, verb, adjectives). German and Turkish syllables were maximally matched (Table 1) with respect to the overall distribution of the manner of articulation for the onset phoneme of each syllable and the percentage of syllabic consonant-vowel CV structure.

**Table 1.** Syllable Transitions.

| **German** |  |  |  |  |
| --- | --- | --- | --- | --- |
| Measurment | Within word | Between word | Difference | p-value |
| Syllable identity | 0.91 | 0.02 | 0.89 | < .0001 |
| Syllable CV | 0.51 | 0.29 | 0.23 | < .01 |
| Onset | 0.36 | 0.05 | 0.31 | < .0001 |
| Onset manner | 0.44 | 0.22 | 0.22 | > .1 |
| Rhyme | 0.51 | 0.05 | 0.46 | < .0001 |

| **Turkish** |  |  |  |  |
| --- | --- | --- | --- | --- |
| Measurment | Within word | Between word | Difference | p-value |
| Syllable identity | 0.75 | 0.02 | 0.73 | < .0001 |
| Syllable CV | 0.57 | 0.34 | 0.23 | < .05 |
| Onset | 0.24 | 0.09 | 0.15 | < .0001 |
| Onset manner | 0.26 | 0.34 | -0.07 | > .1 |
| Rhyme | 0.47 | 0.05 | 0.42 | < .0001 |

*For each phonological measurement average transitional probabilities between consecutive syllables within word boundary and between word boundary in German and Turkish sequences are displayed. Mann-Whitney-Wilcoxon tests revealed significantly higher within word transitional probabilities for all the measurements except for onset manner (where a trend was observed). The presence of phonological patterning at the word (2 Hz) rate in the Turkish sequences, allows for temporal grouping of Turkish syllable pairs by German listeners in the absence of lexical processes.*

#### Stimulus Processing

Syllable stimuli produced by a female German/Turkish bilingual speaker were recorded using an external audio card (Brand) (44100 Hz sampling rate). We recorded individual syllables in isolation (randomly presented to the speaker). The mean duration for German syllables was 358 ms (sd: 72 ms), for Turkish 334 ms (sd: 58 ms). Using this method, we eliminated any co-articulation and prosodic modulation between the two syllables within each word, such that a first syllable contains no acoustic cues that would allow the prediction of the second syllable (Fig. 1 E). Consequently, the prediction of the second syllable relies on higher level linguistic processing (e.g. lexical access of the word and syllable transition information).

Using Praat Vocal Toolkit (http://www.praatvocaltoolkit.com/; Boersma, 2002), the selected syllables tokens were high-pass filtered at 60 Hz, compressed in duration (250 ms; Note that syllables starting with a plosive consonant were compressed to 230 ms and a 20 ms silence period was added in the beginning to simulate the oral occlusion before the burst of plosive consonants), and normalized in peak-amplitude and pitch contour (at 250 Hz). Word stimuli were created by concatenating the two syllables of each word. For the Non-Turkish condition, disyllabic pseudo-word stimuli were created by concatenating two syllables that were quasi-randomly selected from all Turkish syllable stimuli (equal probability of first/second syllable position). Each sequence was created by concatenating randomly selected disyllabic stimuli. For the German conditions, since the word list contained several grammatical classes, we specifically checked the word order of each sequence to eliminate all possible formation of linguistic phrases as well as compound words by consecutive words. For Turkish and Non-Turkish conditions, sequences were checked to avoid ‘German-like’ homophones. Overall, three different sets of sequences were created for each condition.

#### Procedure

Participants were seated in the MEG testing booth in front of a board for instructions. Stimuli were presented binaurally via insert ear-plugs (E-A-RTONE Gold 3A Insert Earphones, Ulrich Keller Medizin-Technik, Weinheim, Germany). Participants’ responses were collected with two button boxes (Current Designs, Inc.). Prior to the experiment, the MEG was recorded while participants listened to a sequence of ‘auditory cortex localizer sounds’ and held their gaze at a fixation cross (pure tones : 0.4 sec tone duration; 250 Hz and 1000 Hz, 100 repetitions, jittered intertrial interval 0-5-1.5 sec). In the experiment, during each trial participants held their gaze at a fixation cross while listening to the auditory sequences. The experiment was run using Psychophysics Toolbox (Brainard, 1997).

Both experiments contained 210 trials (105 per condition). Trials were grouped into 15 blocks. In total, each German, and Turkish disyllabic unit was repeated 15 times. The overall duration for the experiment was 150 minutes, including 90 minutes of recording time and 60 minutes of preparation time, breaks, and post-recording questionnaires.

### MRI and MEG Data Acquisition

The MRI scanning was acquired on a 3 Tesla scanner (Siemens Magnetom Trio, Siemens, Erlangen, Germany). Vitamin-E capsules were used to mark anatomical landmarks (nasion, left and right pre-auricular points). For the MEG recordings, a 269-channel whole-head system (Omega 2000, CTF Systems Inc.) situated in a magnetically shielded room was used. Data recording was performed with a sampling rate of 1200 Hz, an online low pass filtered (cut-off: 300 Hz) and online denoising (higher-order gradiometer balancing). The head position relative to the MEG sensors was continuously tracked and head displacement was corrected in the breaks using the fieldtrip toolbox (http://fieldtrip.fcdonders.nl) (Stolk et al., 2013).

## Statistical Analysis

### Acoustic Analysis

The speech envelope was computed separately for each sentence. The acoustic waveforms were filtered in 8 frequency bands that are equidistant on the cochlear map (between 100 and 8,000 Hz; third-order Butterworth filter; forward and reverse) (Smith et al., 2002). The speech envelop was computed by averaging the magnitude of the Hilbert transformed signal of the 8 frequency bands separately for each sentence. The envelope was resampled to 500 Hz to match the MEG data sampling rate. Next the data was averaged across all trials of a sequence, separately per condition. The mean spectral power across all sequences of an experiment, as well as the standard error across sequences, is plotted separately for the conditions.

### Behavioral Measures Analysis

The analysis is described in the results section.

### MRI Data Analysis

For MRI and MEG data analyses, we used the FieldTrip toolbox (http://fieldtrip.fcdonders.nl) (Oostenveld et al., 2011).

From the individual MRIs of all participants probabilistic tissue maps (including cerebrospinal fluid white and gray matter) were retrieved. In case an individual MRI was missing, the standard Montreal Neurological Institute (MNI) template brain was used. In a next step, the physical relation between sensors and sources was obtained using a single shell volume conduction model (Nolte, 2003). The linear warp transformation was computed between the individual T1 MRI and the MNI template T1. The inverse of that transformation was computed, that is, a template 8 mm grid defined on the MNI template T1 was inversely transformed so that it was warped on the individual head space, based on the individual MRI and the location of the coils during the MEG recording. A leadfield (forward model) was calculated based on the warped MNI grid and the probabilistic tissue map, and used for source reconstruction. This allowed computing statistics across subjects in the MNI space with the grids of all subjects being aligned to each other.

### MEG Data Analysis – Preprocessing

For preprocessing, the data were band-pass filtered off-line (1–160 Hz, Butterworth filter; filter order 4) and line-noise was removed using bandstop filters (49.5-50.5, 99.5-100.5, 149.5-150.5 Hz, two-pass; filter order 4). In a common semi-automatic artifact detection procedure (i.e. the output of the automatic detection was monitored), the signal was filtered in a frequency range that typically contains muscular artifacts (band-pass: 110-140 Hz) or jump artifacts (median filter) and z-normalized per time point and sensor. To accumulate evidence for artifacts that typically occur in more than one sensor, the z-scores were averaged over sensors. We excluded trials exceeding a predefined z-value (muscular artifacts, z = 15; jumps, z = 30). Slow artifacts were removed by rejecting trials were the range (min-max difference) in any channels exceeded a threshold (threshold = 0.75e-5). The data was down-sampled to 500 Hz. And epoched (−2.1 to 9.6 sec). Trials with head movements that exceeded a threshold (5 mm) were rejected. Afterwards, the different blocks of recorded MEG data were concatenated (Note that for each block, during the recording, the head position was adjusted to the initial position of the first block). Sensors with high variance were rejected.

Eye-blink, eye-movement and heartbeat-related artifacts were removed, using independent component analysis (infomax algorithm; Makeig et al., 1996). Components were first reduced to 64 components using principal component analysis. Only in the case of a conclusive conjunction of component topography, time course and variance across trials components were rejected. For the sensor space analysis, spherical spline interpolation was used to interpolate the missing sensors (Perrin et al., 1989).

Trials with correct responses were selected and the trial number was matched between the conditions by randomly selecting trials of the condition with less trials (trial number, Exp. 1: mean = 73.22, SD= 11.02; Exp. 2: mean = 68.68, SD=10.27).

### MEG Data Analysis – Power

For the sensor space analysis, the data was interpolated towards a standard gradiometer location based on the headmodel. It was epoched using a time window of 0.5-9.5 sec (0-0.5 sec after stimulus onset were excluded to avoid onset-related contamination) and averaged across all trials of a condition. Evoked power was computed using singletaper frequency transformation (1-7 Hz) separately for each participant of the two experiments at each condition (frequency resolution: 0.1111 Hz). At each frequency the power was contrasted by the neighbouring frequency bins (+-2-3 bins). Cluster-based permutation tests using Monte-Carlo estimation (Maris and Oostenveld, 2007) were performed to analyze differences between the conditions within each experiment (German vs. Turkish/Non-Turkish; dependent-sample T-statistics) and across experiments (German vs. German and Turkish vs. Non-Turkish; independent-sample T-statistics) with an iteration of the condition affiliation (500 iterations). Adjacent sensor points (minimum number of neighbourhood sensors = 2) with t-values exceeding 97.5% of the permutation distribution were spatially clustered. Clusters were considered significant when the summed t-values with the clusters exceeded 97.5% of the permutation distribution (p < .025; two-sided).

Dynamic Imaging of Coherent Sources (DICS) was used to localize neuronal power (Gross et al., 2001). First, based on the individual leadfields a common source filter (1.3332-4.6662 Hz) was computed across conditions for each participant (lamda = 100%; 0.8 cm grid). Second, based on the filter and fourier transformed data (multi-taper frequency transformation; 0.1111 Hz resolution) the power at 2 Hz and 4 Hz was localized and contrasted with the neighbouring frequency bins (+-2-3 bins). Differences in source power at 2 Hz and 4 Hz were tested using cluster-based permutation tests (1000 iterations; two-sided) to analyze differences between the conditions within each experiment (German vs. Turkish and German vs. Non-Turkish; dependent-sample T-statistics) and across experiments (German vs. German and Turkish vs. Non-Turkish; independent-sample T-statistics) with an iteration of the condition affiliation. Adjacent voxels with t-values exceeding 95% of the permutation distribution were spatially clustered. Clusters were considered significant when the summed t-values with the clusters exceeded 95% of the permutation distribution (two-sided).

The Brainetome atlas (Fan et al., 2016) was used to define ROIs (the left and right STG ROIs: STG1 ROI: A41_42_L/R, STG2 ROI: TE1.0_TE1.2_L/R, MTG ROI: aSTS, SMG ROI: IPL A40rv, IFG ROI: A44v, PCG ROI: A6cdl) to further test the condition differences at 2 Hz revealed in the cluster-test analysis. Differences between conditions at each ROI were tested separately for the hemispheres and the comparisons within each experiment (German vs. Turkish and German vs. Non-Turkish; Wilcoxon signed-rank tests) and across experiments (German vs. German and Turkish vs. Non-Turkish; Mann-Whitney-Wilcoxon test’s). Bonferroni correction across ROIs and hemispheres was applied to correct for inflated p-values.

### MEG Data Analysis – Auditory Localizer

The M100 component elicited by the localizer sounds was analyzed in order to determine sensors that reflect individual auditory cortex activity, which were used as ROI for the sensor space connectivity analysis. After preprocessing, the epoched (0-0.5 sec) trials were baseline corrected (−0.4-0 sec). The M100 peak was determined for each participant, as the max. amplitude of the root mean squared data within a window of 0.07-0.160 sec. The 15 sensors with max. amplitude at this time point were selected separately for each hemisphere and participant. For 5 participants no localizer MEG recordings were present (Exp. 1, n = 4) and thus the M100 sensors were selected based on the grand average of the respective experiment.

### MEG Data Analysis – Connectivity

For the group analysis, the data was interpolated towards a standard gradiometer location based on the headmodel. The Debiased Weighted Phase Lag Index (dwPLI) (Stam et al., 2007; Vinck et al., 2011) was computed at 2 Hz and 4 Hz based on the spectral complex coefficients (singletaper frequency transformation; 0.1111 Hz resolution) of each trial (values between 0-1). The connectivity between the individual M100 sensor ROI (Fig. 4 A, B; Note that for this analysis the localizer data was interpolated prior to the M100 sensor selection) and all other sensors was estimated. Fisher z-transformation was applied to normalize the data prior to further analyzes (for displaying purposes the hyperbolic tangent was applied). The dwPLI values were averaged across all individual M100 sensors. The connectivity of the ROI with itself was set to zero. Differences in sensor space connectivity at 2 Hz and 4 Hz at the M100 ROI were tested using cluster-based permutation tests to analyze differences between the conditions within each experiment (German vs. Turkish and German vs. Non-Turkish; dependent-sample T-statistics) with an iteration of the condition affiliation (500 iterations). Adjacent sensors with t-values exceeding 95% of the permutation distribution were spatially clustered. Clusters were considered significant when the summed t-values with the clusters exceeded 95% of the permutation distribution (one-sided).

**Figure 2.**
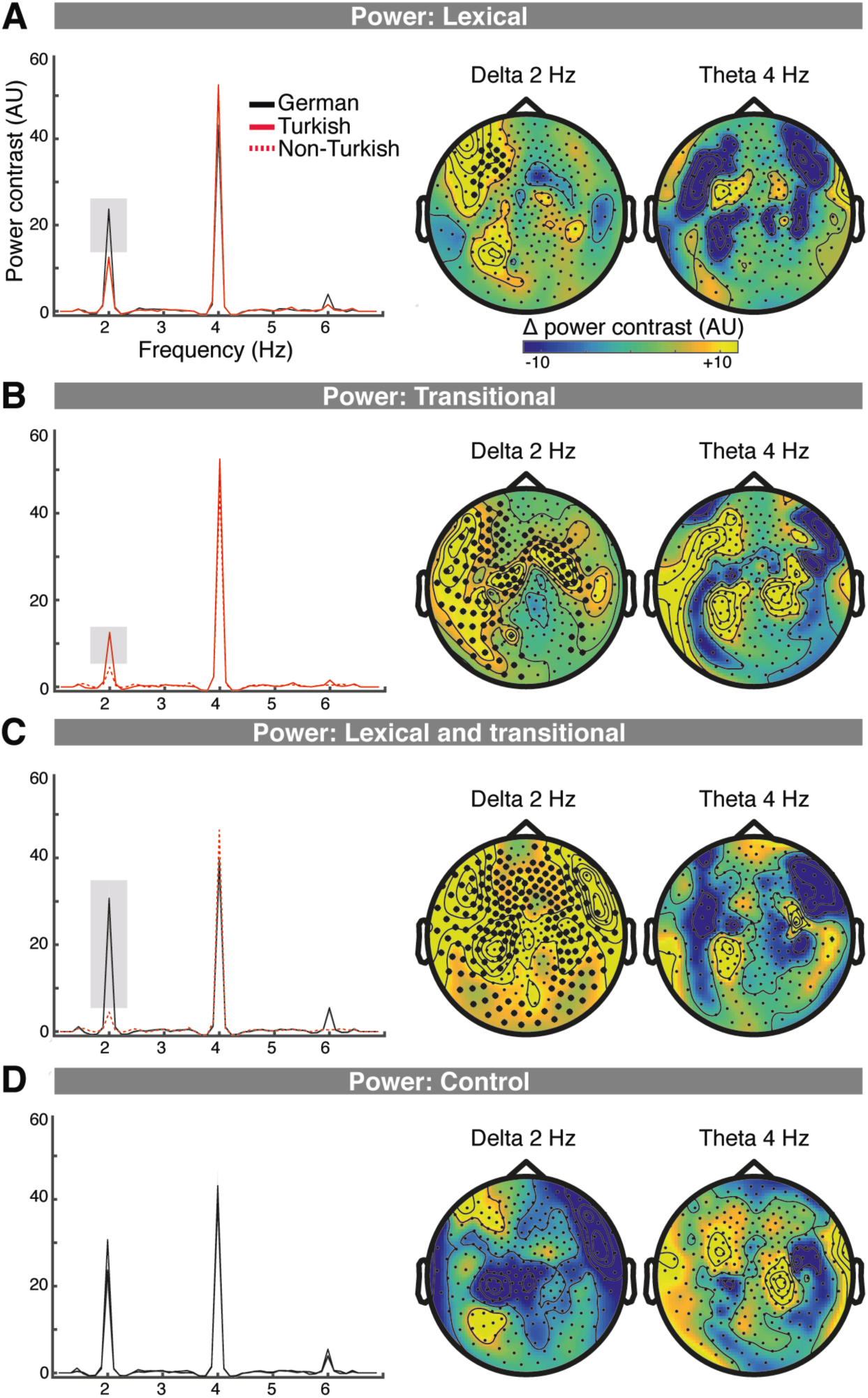
Lexical and Syllable Transition Processing Increases Sensor Space Power. (A-D left column) Power contrasts (with the neighbouring frequency bins) averaged across the individual M100 sensors are displayed; (A-D right column) The topography of the power contrast differences between conditions is displayed at 2 Hz and 4 Hz. Clusters that showed significant differences are marked with black dots; (A) Lexical processing, in Exp. 1, resulted in increased power at 2 Hz for the German compared to the Turkish condition at a left frontal cluster; (B) Syllable transition processing, resulted in increased power in the Turkish compared to the Non-Turkish condition at 2 Hz at a left fronto-central and temporal, and a right fronto-central cluster; (C) Lexical plus syllable transition processing, in Exp. 2, resulted in increased power at 2 Hz in the German compared to the Turkish condition at a broadly distributed cluster; (D) No differences were detected for the across experiment comparison of the German conditions.

**Figure 3.**
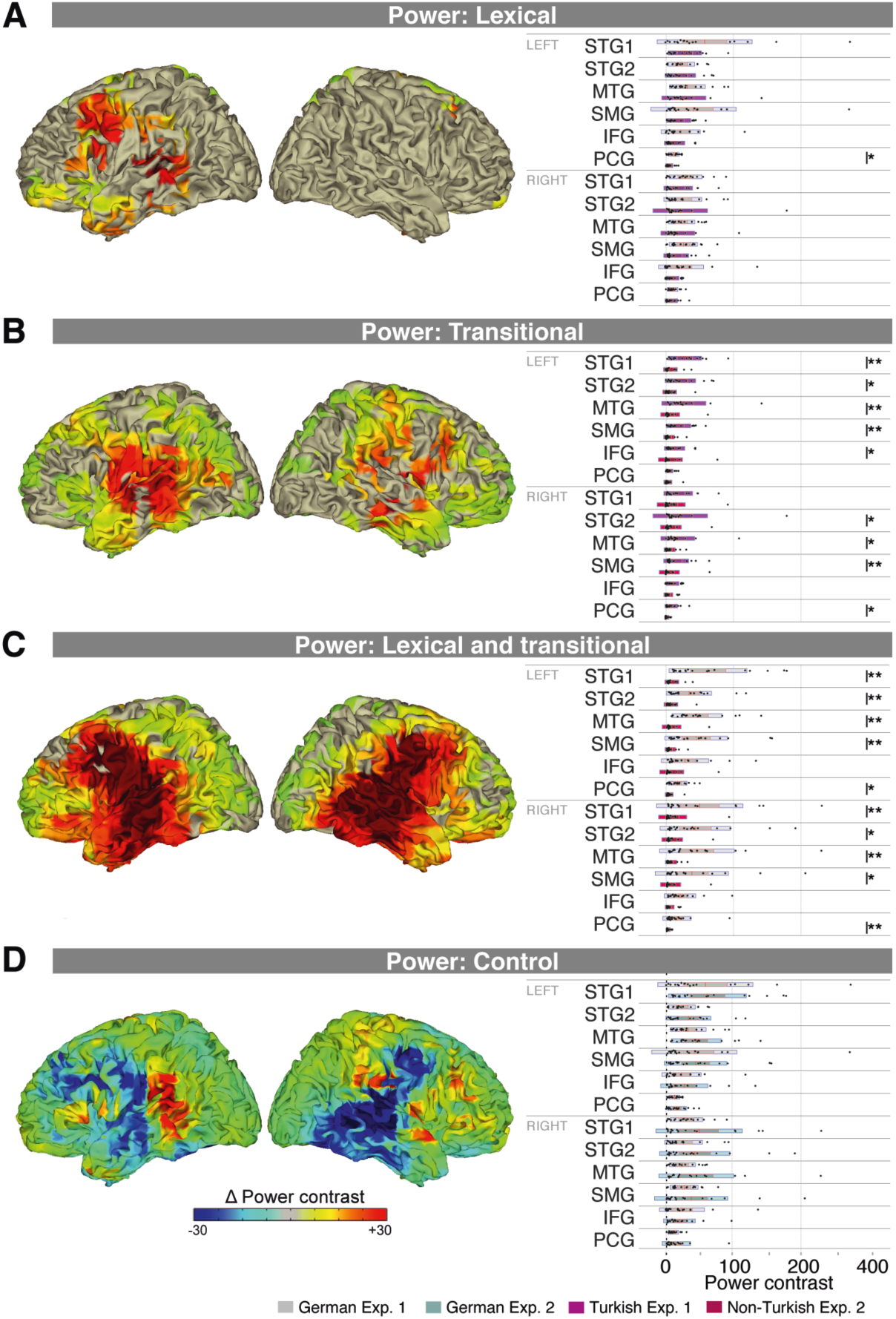
Lexical and Syllable Transition Processing Activates Frontal and Temporal Cortex. (A) (left column) In Exp. 1, lexical processing resulted in increased power at 2 Hz in the German compared to the Turkish condition in two fronto-central-temporal clusters. (right column) Condition differences were significant in the left SMG and there was a trend in the left PMC; (B) (left column) Syllable transition processing resulted in a broad left and a broad right hemispheric cluster showing power increases at 2 Hz; (right column) Condition differences were significant in several left and right hemispheric ROIs; (C) (left column) Lexical plus syllable transition processing resulted in a broad bilateral cluster showing power increases; (right column) Condition differences were significant in several left and right hemispheric ROIs; (D) No significant differences were revealed in the German conditions across experiments;

**Figure 4.**
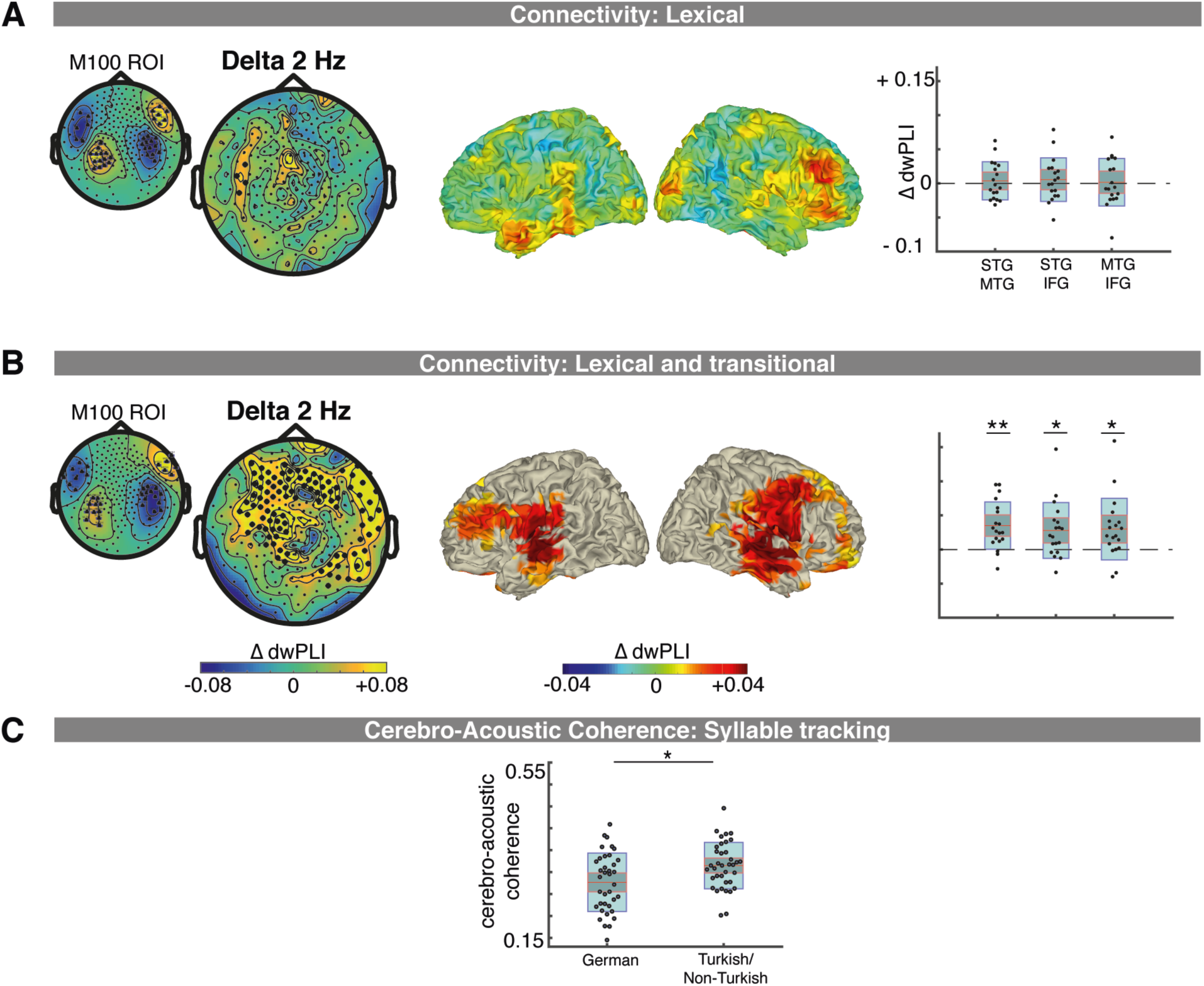
Frontal and Temporal Brain Areas Interact During Lexical and Syllable Transition Processing. (A) Lexical processing: (Left column) The individual M100 sensors were used as region of interest (ROI) for the sensor space connectivity analyses (Exp. 1: the grand average M100 sensors are displayed). In Exp. 1, in the German compared to the Turkish condition connectivity increased at 2 Hz between the M100 ROI and a left fronto-temporal cluster; (Middle column) No significant clusters were detected that showed differences in source connectivity for this comparison (right column) The scatter plot displays condition differences between the STG, MTG, IFG ROI pairs (outer box:1 SD; inner box: 1.96 SEM; red line: mean). (B) Lexical and syllable transitional processing: (Left column) The M100 ROI for Exp. 2 is displayed. In Exp. 2, in the German compared to the Non-Turkish condition connectivity increased at 2 Hz between the M100 ROI and a fronto-temporo-central cluster; (Middle column) The source connectivity between STG and a left/right middle temporal, and fronto-central cluster increased at 2 Hz for the German vs. the Non-Turkish condition; (Right columns) source connectivity was significantly increased between the STG and MTG, STG and IFG and MTG and IFG ROI at 2 Hz. (C) Cerebro-acoustic coherence was significantly reduced in the German compared to the Turkish/Non-Turkish condition.

For the source space connectivity analysis, the spectral complex coefficients were multiplied with a common filter (DICS; across 2 Hz and 4 Hz), computed across conditions separately for each trial. The dwPLI was computed between all voxel. Fisher z-transformation was applied to normalize the data prior to further analyzes. The connectivity between a STG ROI and all voxel was computed by averaging across the voxel of the STG ROIs (A41_42_L/R and TE1.0_TE1.2_L/R). The connectivity of the ROI with itself was set to zero. Source space dwPLI differences between the conditions within each experiment (German vs. Turkish and German vs. Non-Turkish; dependent-sample T-statistics) were tested using cluster-based permutation tests (1000 iterations; see above).

For a more specific analysis of our hypothesis, additionally to the STG ROI, an MTG (aSTS_L/R), and IFG ROI (A44v_L/R) were selected. The connectivity between pairs of ROIs was computed by averaging across the voxels of a ROI and the corresponding ROI pair. Wilcoxon signed-rank and Mann-Whitney-Wilcoxon tests were used to test connectivity differences between conditions separately at each ROI pair. Bonferroni correction was applied to correct for inflated p-values due to multiple-comparisons of ROI-pairs.

Cross-Frequency Coupling: In order to analyze interactions between the 2 Hz word-level and 4 Hz syllable level network, phase-amplitude coupling between the 2 Hz phase and 4 Hz power was computed. After trials were split into 2 seconds long time windows (to increase the amount of data points), the Fourier Transform was used to compute the complex spectral coefficients at 2 Hz and at 4 Hz (0.5 Hz resolution) separately for each trial, condition and participant. A common filter (DICS; lamda = 100%; 0.8 cm grid; 0.5 Hz resolution) was computed at 2 Hz and at 4 Hz and used to project each trial in source space. According to its phase at 2 Hz, the power value at 4 Hz of each trial was sorted into four phase-bins (-pi to pi) separately for each of the ROIs (STG, MTG, IFG), condition and participant. Power was averaged, first across trials, second across the voxels of each ROI, separately for the phase-bins, conditions and subjects. Prior to the statistics each phase-bin was normalized by contrasting it with the average of the remaining phase bins. Nonparametric Wilcoxon Signed-Rank and Mann-Whitney-Wilcoxon tests were used to test differences in power distribution across phase-bins separately for each condition (the data from the German conditions of the two experiments were merged). Bonferroni correction was applied to test for multiple comparisons of phase-bins and ROIs.

Cerebro-Acoustic Coherence: After the spectral complex coefficients at 4 Hz were computed for the speech envelope of each trial (cf. Acoustic Analysis) and the neuronal data (0.1111 Hz resolution), coherence between all sensors and the speech envelope was computed. A common filter (DICS; lamda = 100%; 0.8 cm grid) was multiplied with the coherence, and Fisher z-transformation was applied. The cerebro-acoustic coherence was averaged across voxels of the STG ROIs. Wilcoxon signed-rank tests were used to test the condition difference.

Granger causality: The direction of causal influences between brain areas (i.e. feedforward versus feedback) was examined. Linearly Constrained Minimum Variance (LCMV) beamforming was employed to project and localize the sensor-space measurements in source-space inside the brain volume (Van Veen et al., 1997). First, based on the individual leadfields a common spatial filter was computed across conditions for each participant (lamda = 100%; 0.8 cm grid). Principal component analysis (PCA) (Edward Jackson, 1988) was performed on the time-series of the selected voxels of the ROIs (STG, MTG, IFG ROI) in order to identify the number of principal components that cumulatively explain most of the variance (99.5%). The remaining components were excluded from further analysis. Granger causality analysis was computed by, first fitting the coefficients of a multivariate autoregressive linear model (biosig method; model order: 40) on the time-series, then computing the Fourier transform of these multivariate autoregressive coefficients in the frequency range (1-40 Hz) in order to estimate the transfer function between the model variables, and finally by computing Granger causality from this spectral transfer function. A random condition was generated by shuffling the order of trials separately for each voxel. Fisher z-transformation was applied. Visual inspection revealed peaks in the grandaverage - of the condition differences as well as separately per condition - between 6-8 Hz. For the statistical analysis the Granger causality spectrum was averaged across the voxels of each ROI. Non-parametric paired permutation statistics (1000 iterations; using the MEG & EEG - Toolbox of Hamburg, METH, by Guido Nolte) were used to test the granger causality at 6-8 Hz at each condition against the random condition and between the conditions of each experiment, by randomly permuting the condition affiliation and comparing the observed condition differences against the difference of the randomly permuted cases. Multiple comparisons of ROI contrasts were controlled for using Bonferroni correction for multiple comparisons across ROI pairs (STG-MTG, STG-IFG, MTG-STG, MTG-IFG, IFG-STG, IFG-MTG).

### Additional control analysis

First, in order to access the dynamics of power changes across the experiment, evoked sensor space power was computed ‘blockwise’ (data from 3 consecutive blocks was concatenated in order to gain enough trials) and averaged across all sensors, separately per participant and condition (mean number of trials across blocks (SD), Exp. 1, German: 18 (3.05); Turkish: 17.01 (3.76); Exp. 2, German: 17.20 (3.21); Non-Turkish: 16.70 (4.30)). Non-parametric Friedman tests were used to test differences across blocks separately for each experiment at 2 Hz for the Turkish and Non-Turkish condition (Note, here all responses were analyzed to keep a larger amount of trials). Second, differences in the evoked-responses elicited by the first and second syllable of a disyllabic unit were tested, as possible influence on the phase-amplitude coupling. In each experiment and condition, the RMS of the neuronal sensor space activity at the M100 sensors was computed and baseline corrected (−0.4-0 sec). Trials were averaged separately for the first and second syllables in a sequence (Note that the very first and the very last syllable were discarded to avoid a bias of the activity due to on-/offset effects). Wilcoxon signed-rank tests were used to test differences between the first and second syllable ERP separately for each condition and experiment.

## Results

### Behavioral Measures

A repeated-measures ANOVA tests the effect of lexical and syllable transition cues on target discrimination accuracy (Fig. 1 F) in Exp. 1 and 2 (between-subject factor: experiment; within-subject factor: condition). Accuracy was higher in the German compared to the Turkish/Non-Turkish conditions (*F*(1, 35) = 100.214; *p* < 0.001; η^2^ = 0.374; German: 92 % vs Turkish/Non-Turkish: 83%). There was no main effect of experiment (*F*(1, 35) = 1.794; *p* > .1; η^2^= 0.039) or interaction (*F*(1, 35) = 1.303; *p* > .1; η^2^= 0.008). The results indicate that the presence of lexical cues (in the German conditions) facilitated performance.

### Lexical and Syllable Transition Processing Activate Functional Networks at 2 Hz

In the sensor-space MEG analysis, lexical access effects are reflected in Exp. 1 in power increases in the German compared to the Turkish condition at 2 Hz, at a left frontal and left temporal cluster (p = 0.0040; Fig. 2 A). In source space, the comparison revealed differences at 2 Hz at a left fronto-centro-temporal cluster (p = 0.0040; including STG, MTG, insula, supramarginal gyrus) and a right hemisphere cluster (p = 0.0420; including STG, MTG, MFG, frontal operculum, postcentral gyrus, putamen; Fig. 3 A). Non-parametric comparisons performed at 2 Hz separately for the left and right hemisphere and at the STG1, STG2, MTG, SMG, IFG, PCG ROIs revealed condition differences at the left PCG (p = .0038; Bonferroni corrected alpha = .0042).

The cross-experiment comparison shows sensor-space syllable transition processing effects (Turkish vs. Non-Turkish) within a broad left hemispheric cluster (p = 0.0040) and a broad right hemispheric cluster (p = .0080; Fig. 2 B). In source space, syllable transition processing resulted in increased power in the Turkish compared to the Non-Turkish condition at a bilateral frontal, central and temporal cluster (p = .0020; including the STG, MTG, Heschl’s gyrus, precentral gyrus, supramarginal gyrus; Fig. 3 B). Non-parametric comparisons performed at 2 Hz revealed condition differences at the left STG1, STG2, MTG, SMG, IFG ROI (.0001 < ps < .0034; Bonferroni corrected alpha = .0042). In the right hemisphere condition differences were significant at the STG2, MTG, SMG, PCG ROI (.004 < ps < .0037; alpha = .0042).

Lexical plus sublexical processing, in Exp. 2, resulted in sensor power increases in the German compared to the Non-Turkish condition at a bilateral widespread cluster (p = .0040; Fig. 2 C). In source space, connectivity was increased in the German compared to the Non-Turkish condition at a bilateral frontal, central and temporal cluster at 2 Hz (p = .0020; including the STG, MTG, precentral/postcentral gyrus, frontal inferior operculum; Fig. 3 C). Non-parametric comparisons performed at 2 Hz revealed significant condition differences in the left hemisphere at the STG1, STG2, MTG, SMG and PCG ROI (.0001 < ps < .0015; Bonferroni corrected alpha = .0042). In the right hemisphere condition differences were significant at all ROIs (.0001 < ps < .0022).

There were no significant differences detected at any cluster for the cross-experiment control comparison of the German conditions (sensor space: Fig. 2 D; source space: Fig. 3 D) and no condition differences at any ROI of the two hemispheres (ps > .2545). Likewise, there were no effects at 4 Hz at any comparison in sensor or source space.

### Lexical and Syllable Transition Processing Increase Connectivity Within a 2 Hz Network

In sensor space, lexical access (Exp. 1) resulted in connectivity increases between the M100 ROI and a left fronto-temporal cluster at 2 Hz (p = .0020; Fig. 4 A). Source space analysis revealed no cluster with significant differences in the STG connectivity that would indicate lexical access effects (Exp. 1; p-values > .1439). More specific hypotheses were tested by analyzing connectivity increases between the ROIs of interest (STG, MTG, IFG). In Exp. 1, the connectivity increases between the German and Turkish condition did not reach significance at any ROI pair and frequency (2 Hz: p-values > .7439 and 4 Hz: p-values > .1841; Bonferroni corrected alpha = .0083).

Lexical plus syllable transition processing (Exp. 2), resulted in connectivity increases in the German compared to the Non-Turkish condition between the M100 sensors and at a bilateral widespread cluster (p = .0020; Fig. 4 B). In source space, connectivity increased between STG and a right fronto-centro-temporal network (p = .0100; including right superior temporal, mid temporal, inferior- and mid frontal, supra marginal and precentral areas) and a left lateralized one (p = .0480; including superior- and middle temporal, super marginal areas). A more specific analysis revealed, in Exp. 2, at 2 Hz connectivity was increased between all ROIs (STG-MTG: p = .0010; STG-IFG: p = .0079; MTG-IFG: p = .0070; Bonferroni corrected alpha = .0083; Fig. 4 B), no differences were detected at 4 Hz (p-values > .3760).

No clusters were detected at 4 Hz for any comparison.

### Word-Level Processing at 2 Hz Affects Syllable Processing at 4 Hz

Analyses performed separately for all conditions showed cross-frequency coupling (i.e. the 4 Hz power was computed for 4 different phase-bins of the 2 Hz activity) in conditions with lexical and transitional and syllable information (German), with transitional and syllable information (Turkish), but as expected, not with only-syllable information (Non-Turkish) (Fig. 5). In the German conditions (merged across experiments), power varied across phase-bins 1 and 3 in MTG (p = .0014; Bonferroni corrected alpha = .0028); in the Turkish condition across the phase-bins 2 and 3 in MTG (ps < .0006). There was no significant coupling in the Non-Turkish condition (Bonferroni corrected alpha = .0028; p-values > .0046). The findings suggest cross-frequency coupling was present in all but the Non-Turkish condition, which did not require any form of syllable to word-level grouping (i.e. lexical or syllable transition processing).

**Figure 5.**
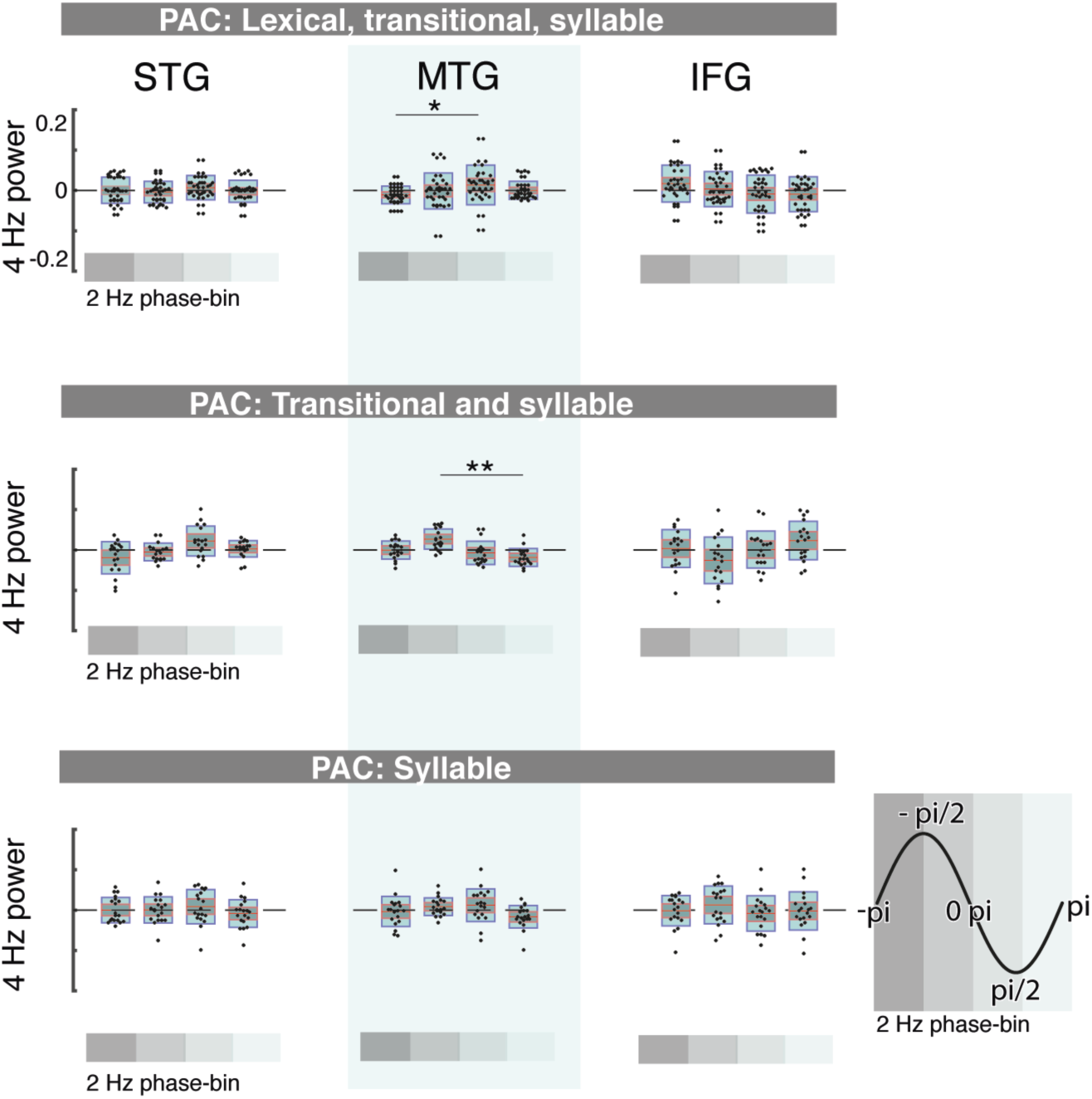
Phase-amplitude coupling at 2 and 4 Hz. The phase-amplitude coupling that is the power at 4 Hz sorted into bins of the 2 Hz phase (legend on the bottom right: gray shades indicate phase-bin 1:4: -pi, -pi/2, 0 pi, pi/2, pi) is displayed for the STG ROI (left column), the MTG ROI (middle column) and the IFG ROI (right column). The phase-amplitude coupling is displayed separately for each condition. First row: Phase-amplitude coupling occurred in the MTG ROI when lexical, transitional and syllable information was present (merged across all German conditions); Second row: it occurred in the MTG ROI when transitional and syllable information was present (Turkish condition); Third row: No coupling occurred when only syllable information was present. Asterixs indicate significant differences between phase-bins. The red line indicates the mean, the dark gray area 1.96 SEM and the light blue area the SD;

An analysis of cerebro-acoustic coherence in STG suggests, interestingly, that word-level processing affects syllable tracking at 4 Hz in STG by decreasing it (Fig. 4 C). As the direction of changes in cerebro-acoustic coherence in the German conditions and Turkish/Non-Turkish conditions were the same, the merged German conditions were tested against the merged Turkish/Non-Turkish conditions, showing significant decreases in the German conditions (p = .0155).

The Granger causality analysis shows significant feedforward and feedback interactions in all conditions when compared to the permuted condition, the STG ROI received feedback from the IFG, but not MTG and provided feedforward input to the MTG and IFG ROI; furthermore MTG and IFG interacted in a feedforward and feedback manner (all p-values < .0083; Bonferroni corrected p-value = .0083). Furthermore, in Exp. 1 in the German condition feedback from IFG to MTG increased compared to the Turkish condition (p = .0040; Bonferroni corrected p-value = .0083). There were no significant effects between the German and Non-Turkish condition in Exp. 2 (p-values > .1120).

### Additional control analyses

First, an additional blockwise analysis on the sensor-space power data revealed in Exp. 1 a tendency towards a blockwise power effect (X^2^(4) = 8.711; p = .069). In Exp. 2, there was no significant effect (X^2^(4) = 5.347; p = .253). However, the post hoc Mann-Whitney-Wilcoxon tests show no significant differences between blocks in any experiment. There were no significant effects at 4 Hz. Second, differences in the ERPs elicited by first and second syllables were analyzed as possible influence on the phase-amplitude coupling. The analysis suggests that the syllable and word-level processing does not reflect differences of the ERPs of the first and second syllable of dysyllabic units (No significant differences were observed; all p-values > .05).

## Discussion

We show that the frequency-tagging paradigm can be used to tease apart aspects of lexical-level and syllable-transition information processing by differentiating neuronal networks activated at 2 Hz (Fig. 6). Speech comprehension models have focused on mapping of acoustic-phonemic to lexical processing, e.g. (Marslen-Wilson and Welsh, 1978), supported by important cognitive neuroscience evidence for the processing and encoding of phonemic level information (Mesgarani et al., 2014; Di Liberto et al., 2015). Other models have suggested, however, that suprasegmental information such as stress or syllable rhythm can facilitate lexical access (Mehler et al., 1981; Cutler, 2012). The data we provide enrich our understanding by implicating contributions of an acoustic-syllabic to lexical processing route. Specifically, we show interactions between lexical and syllable-transition processing of words (here at 2 Hz) and acoustic-syllable processing (here at 4 Hz), first, in terms of phase-amplitude coupling, and second, in terms of a decreased tracking of syllables (cerebro-acoustic coherence; here at 4 Hz) in STG when lexical or syllable transition word-level content was present. At both the syllable level and the word level we are not committed to any decoding scheme. At the syllable level, we show that syllabic information - as a whole unit or as a sequence of phonemes - is obtained within a window-duration that is inside the theta range. The strongest evidence that this window is determined by theta-band oscillations comes from earlier work on the association of the drop in intelligibility of speeded speech with the upper frequency range of theta (Ghitza and Greenberg, 2009; Doelling et al., 2014). At the word level, we do not link our findings on the lexical processing to oscillations.

**Figure 6.**
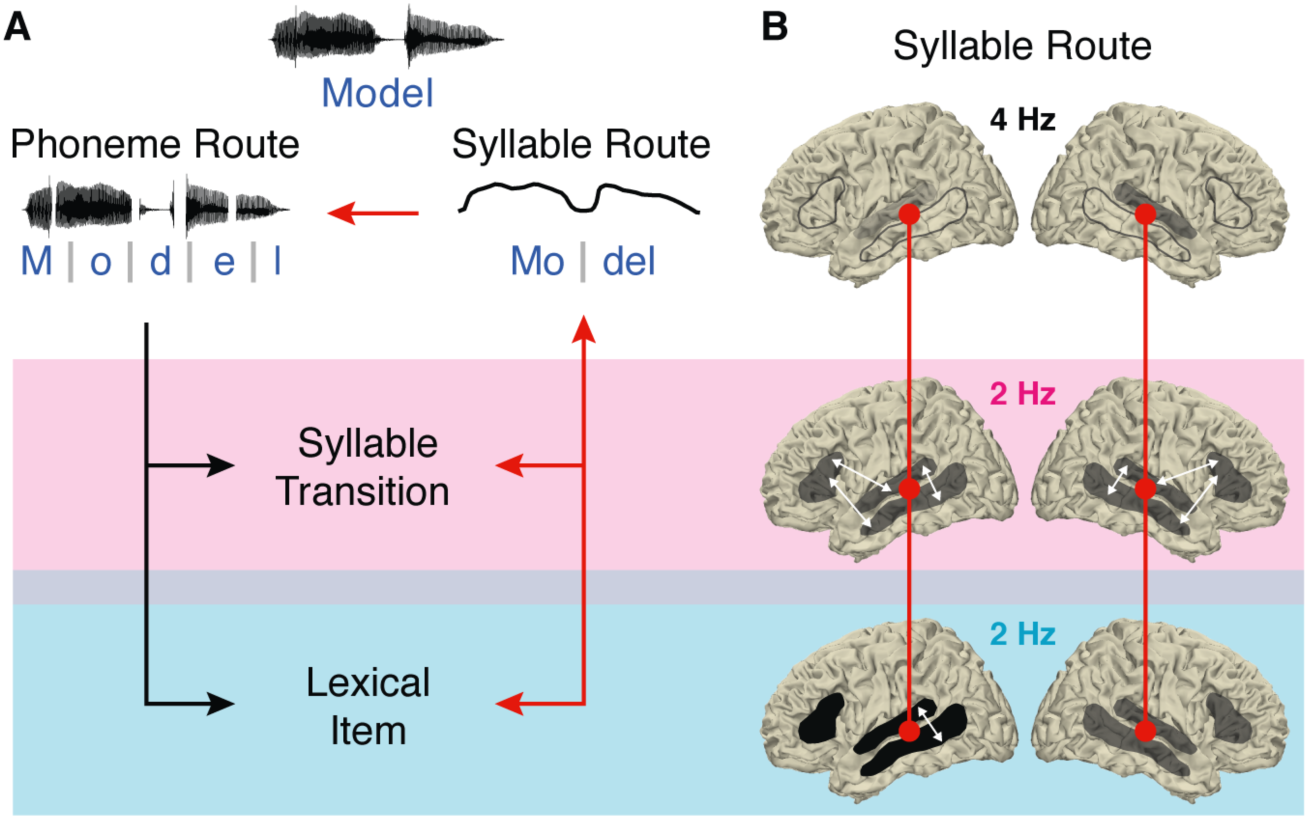
Schematic of the Interaction of Syllable and Word-Level Processing. (A) Extending classic ‘Phoneme Route’ models, we propose contributions of a ‘Syllabic Route’ to word-level processing. (B) (first row) Low-frequency tracking of the syllabic rate in auditory cortex including STG (for 4 Hz power see Fig. S2) affects (second row) higher level syllable transition processing in a network including bilateral STG, MTG and IFG (2 Hz activity) and (third row) lexical processing in a network including STG and left MTG and IFG and interactions between STG and left middle temporal areas (2 Hz activity). The interactions between syllable and word-level processing occur in STG and MTG (red line). (B, C, D) The intensity of the gray shades indicates the activation level during the respective process.

### Lexical and Syllable Transition Processing of Words Activate Networks at 2 Hz

Consistent with previous research, lexical processing activated left fronto-central and middle temporal brain areas, which have been argued to support lexical-semantic processing of words (Fig 2A, 3A) (Hickok and Poeppel, 2004, 2007; Peelle, 2012; Gow, 2012; Rice et al., 2018). ‘Mere’ syllable transition processing, in contrast, activated fronto-centro-temporal brain areas in both hemispheres (Fig 2B, 3B). Previously, a functional subdivision of the temporal cortex has been proposed, with bilateral STS activations during lower level acoustic speech processing and a left-lateralized activation of the more ventral temporal-parietal cortex during lexical-semantic processing (Binder et al., 2000, 2009). In line with this subdivision, our findings further suggest that, beyond acoustic processing, sublexical syllable-transition processing occurs bilaterally. The functional connectivity increases between left middle temporal brain areas and auditory cortex during lexical processing of words, might indicate interactions of higher-level word processing in MTG with lower-level phonological or acoustic processing of words, in line with the view of the pMTG as hub for the mapping between words and meaning that links acoustic phonetic representations in the bilateral pSTG to semantic representations (Kertesz et al., 1982; Gow and Segawa, 2009; Hickok and Poeppel, 2007; Gow, 2012). The bilateral interactions of STG, MTG and IFG when additionally syllable transition information was present, possibly indicate an increased recruitment of articulation or motor prediction related brain areas (as previously activated during sublexical processing) (Hickok and Poeppel, 2000, 2007; Gow, 2012). In our paradigm, increased neuronal activity in the native language condition, which contained semantic and syllable transition cues to group syllables into words, compared to a foreign language condition, which contained only syllable transition cues, indicates lexical processing of words. Lexical processing and syllable transition processing, however, are tightly entangled, thus an alternative possibility is that the observed increase in neuronal activity partly reflects better-learned syllable transitions in a native compared to a foreign language condition.

### Word-Level Networks at 2 Hz and Syllable Networks at 4 Hz Interact

Previous speech comprehension models have focused on mapping of acoustic-phonemic to lexical processing (e.g. (Marslen-Wilson and Welsh, 1978). Neurophysiological data, however, provide compelling evidence for the extraction of information at the syllable level (Gross et al., 2001; Luo and Poeppel, 2007; Panzeri et al., 2010). What does that mean for our understanding of speech comprehension? We provide evidence for interactions between word-level and syllable level processing (phase-amplitude coupling, Fig. 5, 6, and decreased cerebro-acoustic coherence when word information is present), suggesting that syllable information is not only extracted to inform syllable transitions but exploited for lexical-level processing. Word- and syllable-level processing interacted in the MTG during lexical and lexical-plus-syllable transition processing, not however when only syllable content was present (Fig. 6).

The findings suggest, in line with proposals of a crucial role of the MTG as an interface between phonetic and semantic representations (Gow, 2012), that additionally to the STG, the MTG is involved in communicating information between syllable and word-level processing. It is likely that our findings indicate both feedforward communication from STG to higher-level processing areas and feedback from the word-level to the syllable-level (cf. Granger Causality), for example the first syllable might provide (temporal and/or semantic) predictions of the second syllable. Note that the interaction effect we observed cannot be explained by mere differences in the ERPs elicited by the first and second syllables (Fig. S3). Interactions between lexical and phonological processing have been shown to involve feedback from pMTG to pSTG (Gow and Segawa, 2009, for review: Gow et al., 2008). Furthermore, several electrophysiological studies suggest interactions/feedback from sentential (Gow and Olson, 2016) or phrasal processing (Keitel et al., 2018), or possibly both (Park et al., 2015) to syllable processing. However, research that is particularly designed to investigate the interactions at the word level is rare (Mai et al., 2016; Gow and Olson, 2016; Keitel et al., 2018). The frequency-tagging paradigm that we used might be particularly sensitive to revealing interaction effects by generating a higher consistency in the frequency of word processing. Additionally, our findings of decreased speech tracking at 4 Hz in STG (i.e. cerebro-acoustic coherence) when word-level content is present indicate an interaction between the word-level and syllable-level processing and support the notion of feedback communication. While several studies found increased speech tracking during the processing of intelligible compared to unintelligible speech (Peelle et al., 2013; Rimmele et al., 2015; Park et al., 2015), a previous artificial word learning study revealed reduced power elicited by syllable processing when words were intelligible (Buiatti et al., 2009; Sohoglu and Davis, 2016), suggesting that the effects of the interaction can vary depending on the paradigm, performed process etc. This, too, will need to be assessed in further experiments.

### Processing of Syllable Transition Cues

Behavioral research suggests that sequencing of phonemes - because the distribution of phonemes varies across syllables - can be used to detect syllable transitions and word boundaries (McQueen, 1998), as well as the position of syllables within words (van der Lugt, 2001; Cutler, 2012). We showed the neuronal networks involved in using syllable transition information to process disyllabic words (Fig. 2 B, 3 B) and their interaction with syllable-level processing (Fig. 5). Our findings provide evidence that even the syllable transition information present in a foreign language (Table 1), that is, cues such as the onset of a syllable or the consonant-vowel pattern, can be extracted and used to group syllables into words. In the present study, the stimuli were recorded and preprocessed so that acoustical cues at the word level were minimized, resulting in a prominent power peak only at the syllable rate (Fig. 1 E). Thus, the increased power peak at 2 Hz (in the Turkish compared to the Non-Turkish) condition reflects the processing of syllable-transition features rather than the processing of acoustic cues.

Statistical learning of the contingencies throughout the experiment might have additionally contributed to our findings, as the power peak at 2 Hz showed a tendency to vary across consecutive blocks of the experiment. The rapid statistical learning of such cues would be in line with previous studies (Saffran et al., 1996; Pena and Melloni, 2012; Ota and Skarabela, 2016, 2018). Our findings raise new questions. In the current study, we carefully matched the German and Turkish stimulus material with regard to sublexical cues that can be used to group syllables into words. Possibly this enhanced the ability of participants to quickly extract (and possibly learn) the sublexical contingencies of a foreign language. If the ability to extract sublexical contingencies at the word level depends on the similarity of these features between languages, the frequency-tagging paradigm could be used as a neurophysiological tool to investigate the phonological similarity between languages, without requiring explicit feedback from participants.

## Conclusions

Our study provides novel evidence for the interaction of syllable and word-level functional networks. The increased coupling between word- and syllable-level processing, when lexical or syllable transition cues are present, suggests that these processes might be more interactive than previously thought.

## Acknowledgements

This work was funded by the Max-Planck-Institute for Empirical Aesthetics. We thank Marius Schneider for help with the data recording, Ilkay Isik for checking the Turkish stimulus material, Dr. Florencia Assaneo for discussions and Felix Bernoully for graphics support.

